# Sickle cell disease complications: prevalence and resource utilization

**DOI:** 10.1101/577189

**Authors:** Nirmish Shah, Menaka Bhor, Lin Xie, Jincy Paulose, Huseyin Yuce

## Abstract

This study evaluated the prevalence rate of vaso-occlusive crisis (VOC) episodes, rates of uncomplicated and complicated VOC episodes, and the primary reasons for emergency room (ER) visits and inpatient admissions for sickle cell disease (SCD) patients. The Medicaid Analytic extracts database was used to identify adult SCD patients using claims from 01JUL2009-31DEC2012. The date of the first observed SCD claim was designated as the index date. Patients were required to have continuous medical and pharmacy benefits for ≥6 months baseline and ≥12 months follow-up period. Patient demographics, baseline clinical characteristics, the rate of uncomplicated and complicated VOC (VOC with concomitant SCD complications) episodes, and reasons for ER visits and inpatient stays were analyzed descriptively. A total of 8,521 patients were included in the analysis, with a median age of 30 years. The average follow-up period was 2.7 years. The rate of VOC episodes anytime in the follow-up was 3.31 in person-years. During the first-year follow-up period, an average of 2.79 VOC episodes were identified per SCD patients, with 1.06 VOC episodes treated in inpatient setting and 0.90 VOC episodes in ER without admission. A total of 76,154 VOC episodes were identified during the entire follow-up period for the overall SCD patients. Most of the VOC episodes (70.3% [n=53,523]) were uncomplicated episodes, and 29.7% were complicated episodes. Using primary diagnosis claims only, the most frequent complications during the VOC episode were infectious diseases (25.9%), fever (21.8%), and pulmonary disorders (16.2%). Among ER and hospitalizations related to VOC or SCD complication, ~85.0% had VOCs as the primary reason for admission; 15.0% had SCD complications as the primary reason. In summary, SCD and its related comorbidities and complications result in high acute health care utilization. In addition, VOC remains the primary reason for SCD patients’ ER visits and inpatient admissions.

## Introduction

Sickle cell disease (SCD) is a life-threatening genetic disorder affects ~100,000 individuals in the United States, where it is one of the most common genetic blood disorders [1,2]. Damage to the red blood cells (RBCs) occurs due to polymerization of deoxygenated HbS and these damaged cells have abnormal structures as well as expression of adhesion molecules. This results in hemolytic anemia and the blockade of small blood vessels, which lead to vaso-occlusion and end organ failure [3]. Importantly, multicell adhesion between RBCs, white blood cells, platelets, and endothelial cells result in painful vaso-occlusive crisis [4–6]. There are several common subtypes of SCD such as homozygous hemoglobin S (or sickle cell anemia), hemoglobin C, and hemoglobin S/beta-thalassemia [7,8]. SCD

The most common sequelae of SCD include VOC episodes, invasive infections, acute chest syndrome, strokes, and chronic pulmonary hypertension [9]. SCD causes recurrent VOC episodes as well as increases the risk of infections and other complications that often require emergency intervention [10]. VOC is a very important complication of SCD: the higher the number of VOCs experienced by SCD patients, the worse the patient’s quality of life [11,12]. Multicell adhesion is the driver of VOC, and targeting this adhesion results in more VOC-free days [13]. A study that used the National Hospital Ambulatory Medical Care Survey for 1999–2007 showed the most common patient-cited reasons for ER visits were chest pain, other or unspecified pain, fever/infection, and respiratory disorders [14]. Studies have shown that VOC is the main cause of most SCD health care utilization or medical contact, and high health care utilization among SCD patients remains prevalent due to poor acute pain management [15–17]. Furthermore, SCD has been associated with high utilization of medical care, such as ER visits and hospitalization, as observed in a study finding that as high as 53% of SCD patients who had ER visits subsequently required inpatient stays in hospitals or other treatment facilities [18]. A study conducted with Florida Medicaid program data from 2001-2005 across all patients showed that during the 5-year study period, SCD patients incurred an average of 3.7 inpatient hospitalizations and 24.1 hospital days, with ~84% attributable to SCD-related diagnoses [19].

Other studies have found that higher health care utilization in adult SCD patients is associated with a higher risk for mortality, and acute SCD pain episodes are the most commonly observed cause of inpatient admission in SCD patients [20,21].

Although some Medicaid studies have looked at the reason for admissions and ER visits in SCD patients and the rate of pain crisis episodes, very few studies have looked at the rate of SCD complications in different settings using real-world data. Therefore, using real-world data to shed light on the more current complications and reasons for health care utilization in adult SCD patients, this study descriptively evaluates the prevalence rate of VOC, the rate of complicated VOC (VOC with concomitant SCD complications) and uncomplicated VOC episodes, and the primary reasons for ER and inpatient admissions, with the most recent Medicaid database data available.

## Materials and Methods

### Data Sources

This was a retrospective, descriptive study using US Medicaid population data from 01JAN2009-31DEC2013. The MAX data system contains extensive individual-level information on the characteristics of Medicaid enrollees in all 50 states and the District of Columbia as well as the services used during a calendar year. Specifically, MAX consists of 1 personal summary file and 4 claims files that provide fee-for-service claims, managed care encounter data, and premium payments.

The study included fee-for-service patients from all available states and Managed Care enrollees who resided in 14 states that had relatively complete data (Arizona, California, Indiana, Kansas, Kentucky, Minnesota, Nebraska, New Jersey, New Mexico, New York, Oregon, Tennessee, Texas, and Virginia). Service use among Managed Care enrollees is captured in encounter data (ie, data collected from patients when they receive services). Patients who had dual eligibility with Medicare were not included in this study due to the incomplete information in the MAX database.

### Patient Selection

Adult patients were selected if they had ≥1 diagnosed claim with SCD (ICD-9-CM code 282.41-282.42, 282.60-282.69) during the identification period (01JUL2009-31DEC2012). The date of the first observed SCD diagnosis claim was designated as the index date. Patients were required to have continuous health plan enrollment with medical and pharmacy benefits for ≥6 months preindex date (baseline period) and ≥12 months post-index date (follow-up period). Patients were excluded if they were enrolled in a clinical trial during the study period (identified using ICD-9 codes V70.7).

### Baseline Measures

Patient socio-demographic and clinical characteristics were measured, including age, sex, race/ethnicity, geographical region, and Charlson comorbidity index score. In addition, the rate of inpatient stays and ER visits during the baseline period was assessed.

### Outcome Measures

The rate of VOC episodes in person-years during the entire follow-up period and the annual counts of VOC episodes (in total and by place of service in the first year) were assessed. VOC episodes were defined using the following ICD-9 codes (282.42, 282.62, 282.64, 282.69) from both primary and secondary claim positions. VOC claims without a 3-day gap were combined and considered 1 VOC episode. The number of VOC episodes in different settings were also calculated based on place of service using a hierarchical order: inpatient, ER, outpatient, office, and other.

Among the total VOC episodes identified in the follow-up period, the percentage of complicated and uncomplicated VOC episodes were examined [22]. A complicated VOC episode was defined as the presence of a diagnosis of other SCD complications (S1 Table) during the VOC episode, and uncomplicated VOC was defined as no occurrence of any other SCD complication during the VOC episode. The top 10 concomitant SCD complications (using both primary and secondary diagnosis claim positions) were identified.

In addition, the total number of ER and hospital visits related to VOC and SCD complications were examined during the 12-month follow-up period. The primary reasons for SCD-related ER and hospital visits were assessed based on the primary discharge diagnosis codes, including VOC (ICD-9-CM: 282.41, 282.42, 282.60-282.69) and SCD complications (S1 Table). Percentage of ER and inpatient visits associated with VOC or SCD complications were calculated.

### Statistical Methods

All variables were analyzed descriptively. Percentages and numbers were provided for dichotomous and polychotomous variables. Means and standard deviations were examined for continuous variables. For length of stay (LOS) of inpatient admission associated with VOC or SCD complication, median and interquartile range for LOS was also evaluated.

## Results

### Baseline Characteristics

A total of 8,521 patients met the study selection criteria and were included in the analysis (Fig 1). The mean age of patients was 32.88 years (Standard deviation [SD]=12.21), and the median age was 30 years. Patients were mainly from the Northeast region (44.6%). Most patients were female (67.3%) and African American (74%), and the remaining 26% patients were white, Hispanic, other, or unknown. The mean Charlson comorbidity index score was 0.65, and the most frequent comorbid conditions during the baseline period were infectious diseases (19.7%), asthma (10.9%), fever (9.3%), neoplasms (benign and malignant) (7.2%), upper respiratory tract infections (6.6%), and constipation (5.9%).

**Fig 1.**
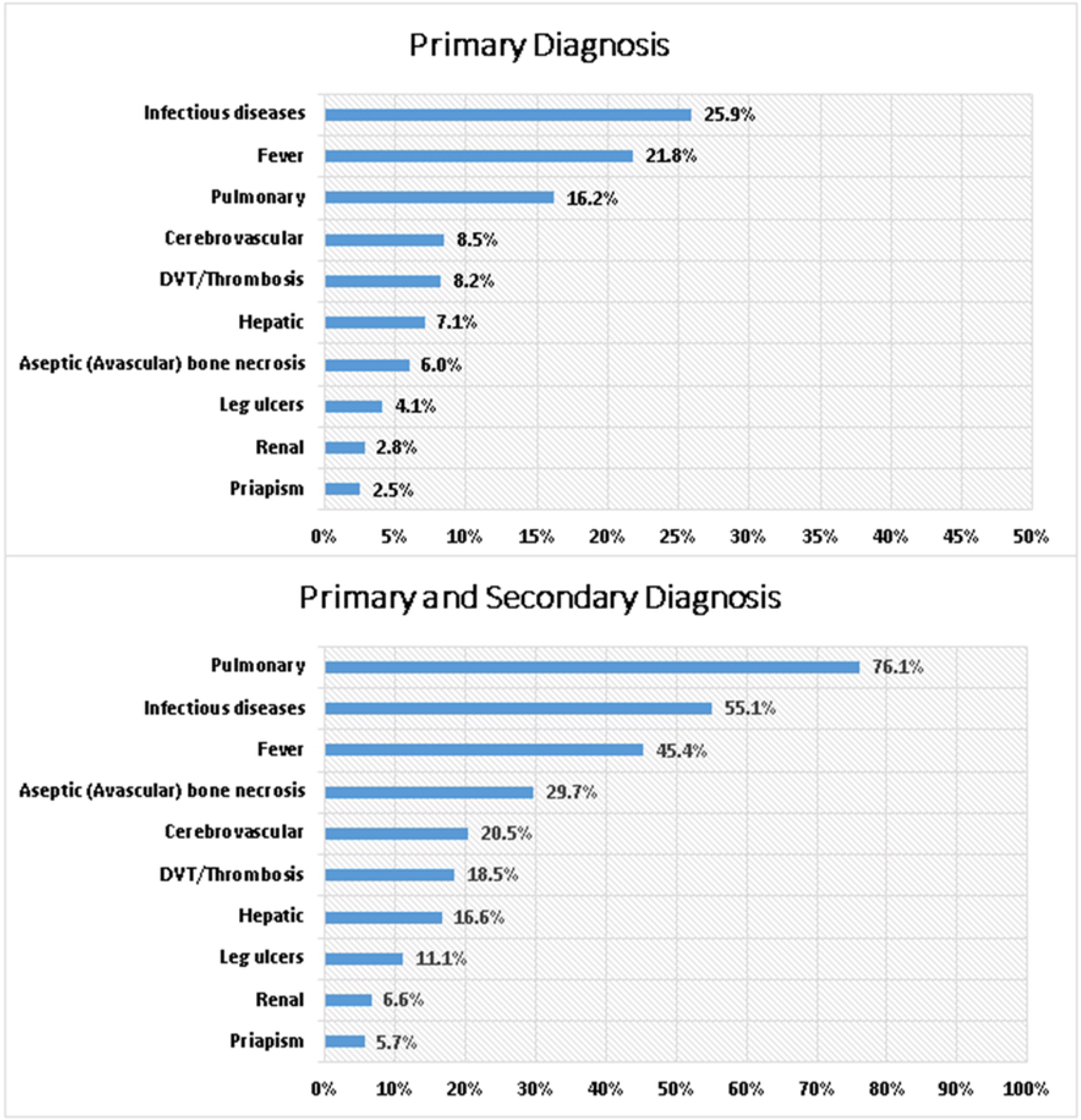
Flow Chart for Patients Selection Criteria. SCD: Sickle cell disease.

During the baseline period, 53.6% (n=4,563) of the SCD patients had ≥1 ER visit and 29.8% (n=2,543) had ≥1 inpatient admission (Table 1).

**Table 1.**
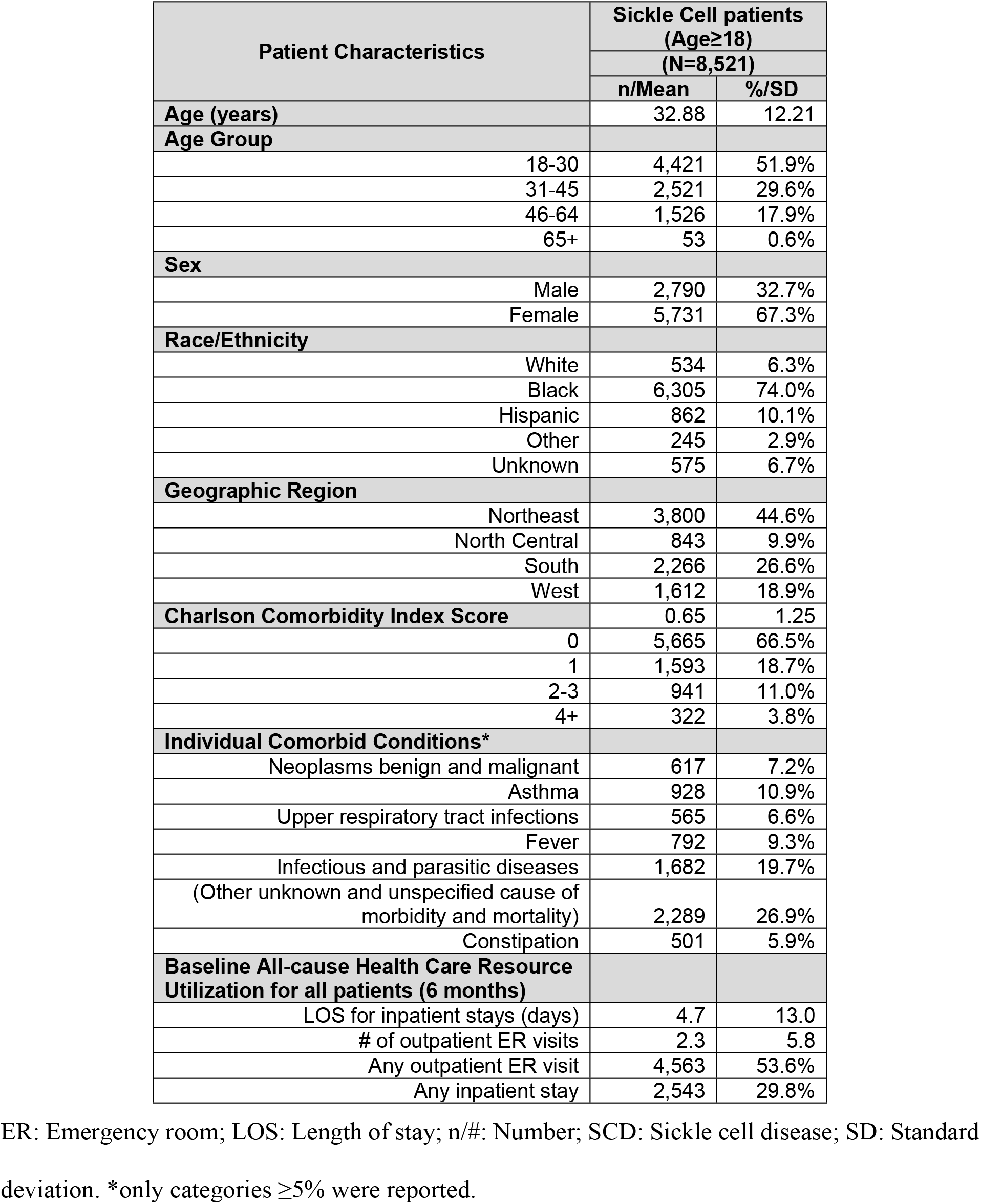
Baseline Demographic and Clinical Characteristics.

## Follow-up Results

### Rate of pain crisis episodes

The mean/(SD) and median length of the follow-up periods were 972/(378) days and 981 days, respectively. During the entire follow-up, 53.4% of the 8,521 patients had ≥ 1 VOC episode, with a rate of 3.31 VOCs per person-year. During the first year of the follow-up period, 52.3% of the patients did not have any VOC episode, 14.7% had 1 VOC episode, 6.7% had 2 VOC episodes and remaining 26.3% had more than 2 VOC episodes (Fig 2). An average of 2.79 VOCs were identified per SCD patient during the first year of follow-up. Out of the total 2.79 VOCs identified per SCD patient, 1.06 VOC episodes were in inpatient setting, 0.90 VOC episodes were in an ER setting, 0.51 VOC episodes were in an outpatient setting, 0.24 VOC episodes were in an office setting, and 0.09 VOC episodes were in another setting such as pharmacy, school, correctional facilities, etc (Fig 3).

**Fig 2.**
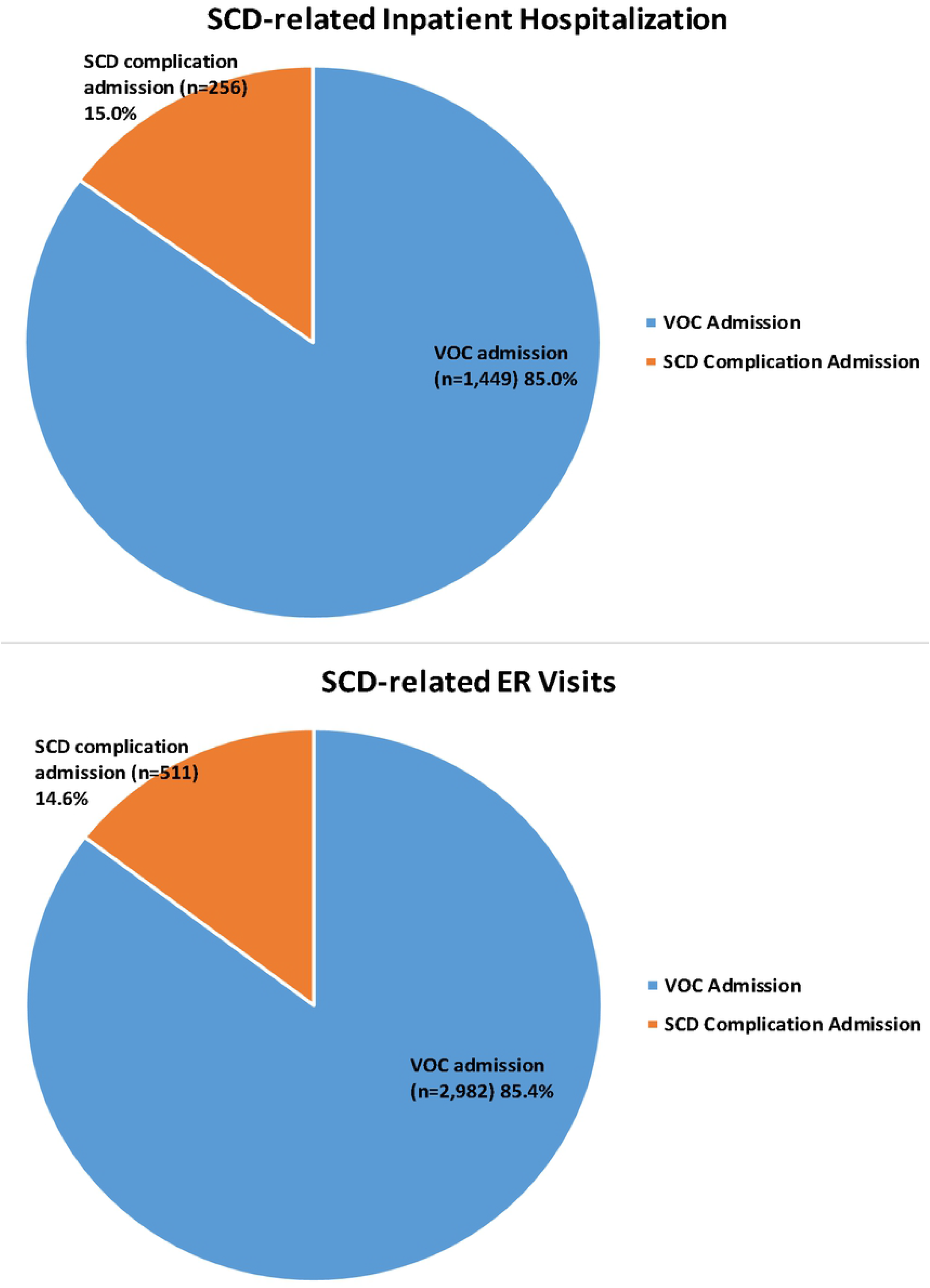
The Frequency of VOC During the First Year of Follow-up. VOC: Vaso-occlusive crisis.

**Fig 3.**
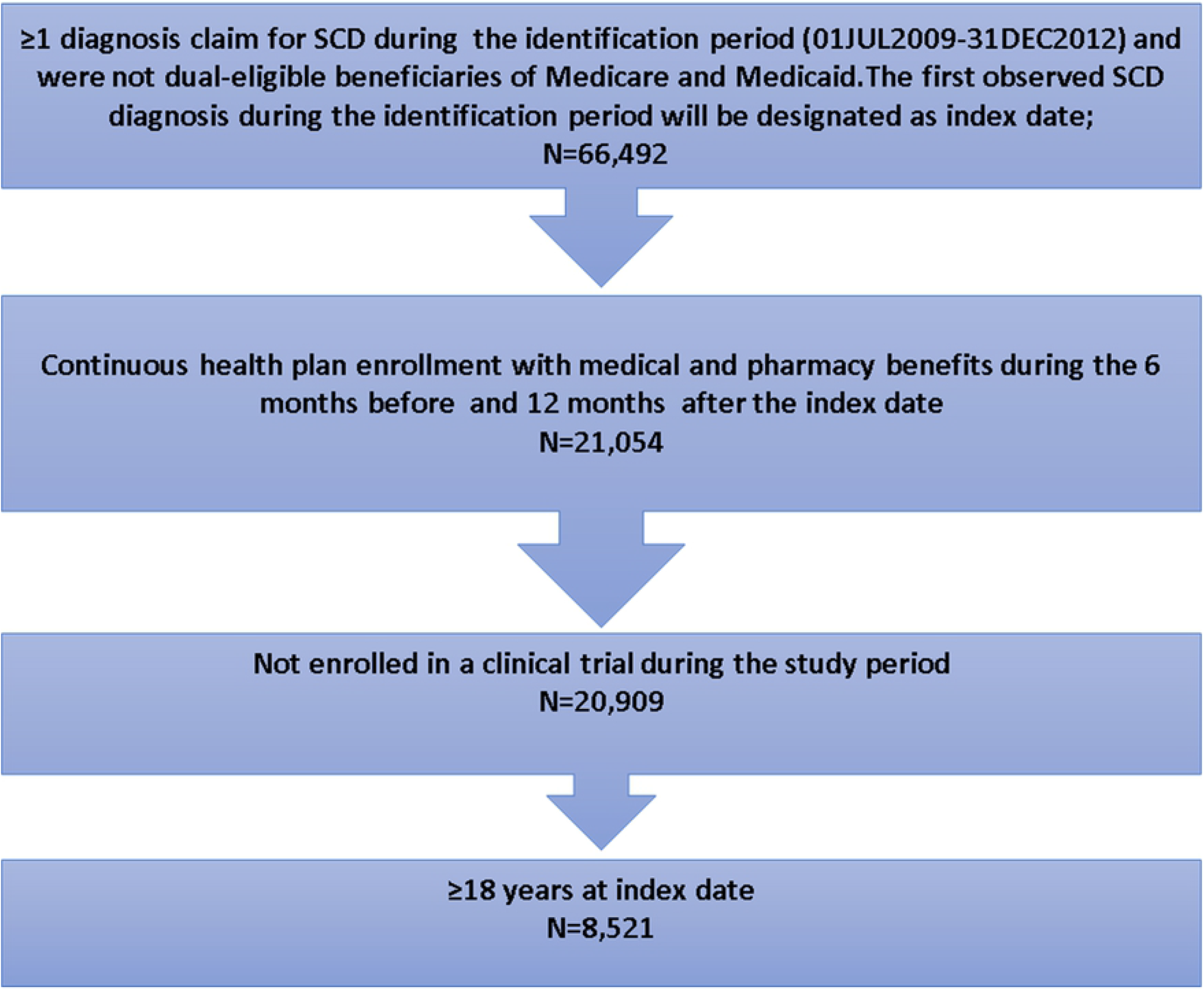
VOC Per Patients in Different Settings During the First Year of Follow-up. ER: Emergency room; VOC: Vaso-occlusive crisis; #: Number; Other setting includes pharmacy, school, homeless shelter, prison/correctional facility, patient’s home, assisted living facility, independent laboratory and unknown setting.

### Rate of Complicated and Uncomplicated VOC episodes

During the entire follow-up period, 76,154 VOC episodes were identified for the 8,521 adult patients. Of the 76,154 VOC episodes observed, 22,631 (29.7%) were complicated VOCs and 53,523 (70.3%) were uncomplicated VOCs, respectively. Of the 22,631 complicated VOC episodes, the top associated concomitant SCD complications (according to primary diagnosis claims only) included infectious diseases (25.9%), fever (21.8%), pulmonary disorders (16.2%), cerebrovascular conditions (8.5%), and thrombosis/deep vein thrombosis (8.2%). While the top associated concomitant SCD complications (according to both primary and secondary diagnosis claim positions) were pulmonary disorders (76.1%), infectious diseases (55.1%), fever (45.4%), aseptic (avascular) bone necrosis (29.7%), and cerebrovascular conditions (20.5%) (Fig 4).

**Fig 4.**
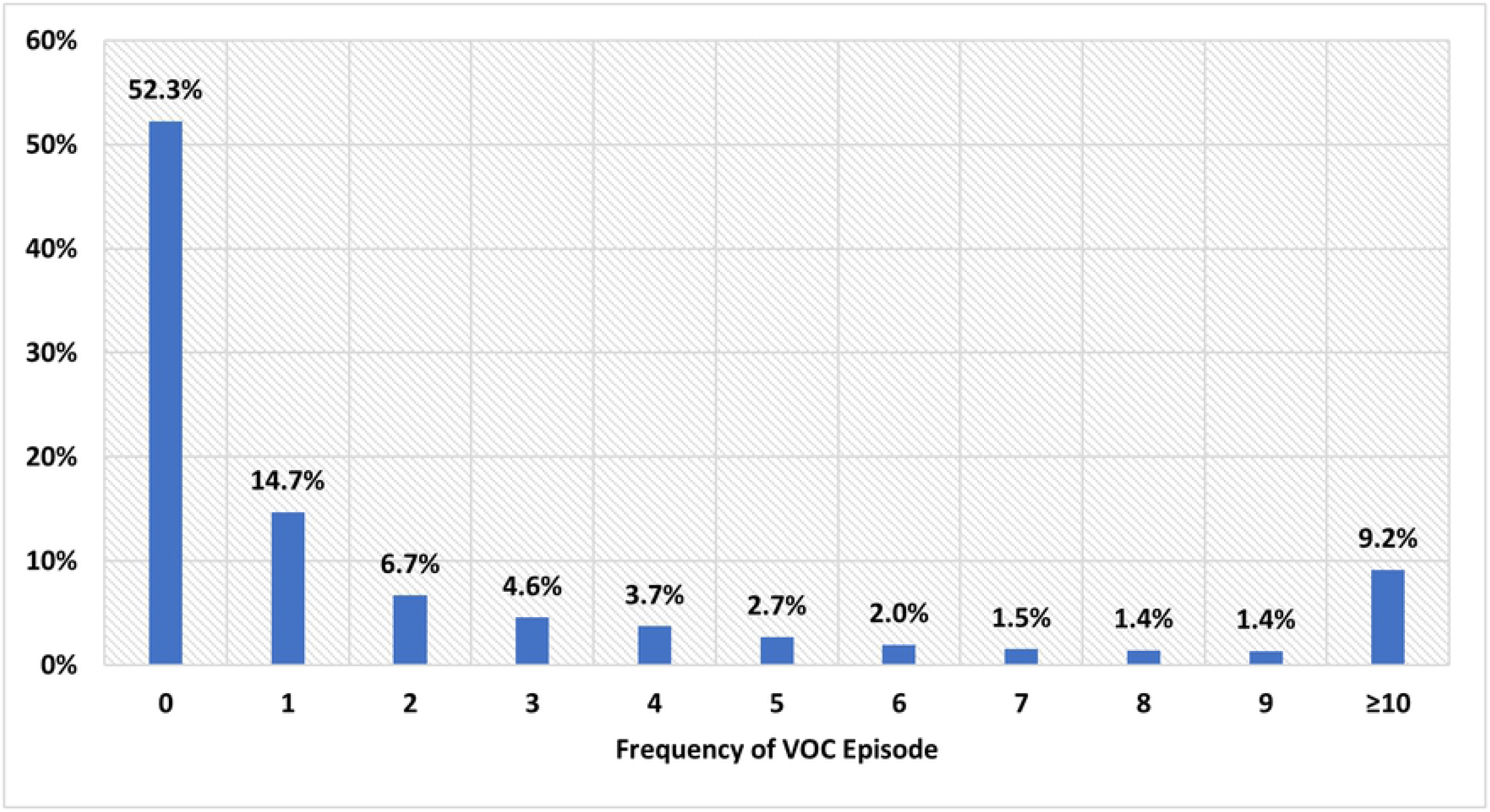
SCD-Related Complications and Concomitant Comorbidities Using Primary and Secondary Diagnosis During the Entire Follow-up Period. DVT: Deep vein thrombosis; SCD: Sickle cell disease.

### Primary reasons for SCD-related ER visits and hospitalization

Of the 8,521 adult SCD patients included in the study, a total of 3,493 ER visits related to VOC or SCD complications were identified during the 1-year follow-up period. Similarly, a total of 1,705 hospital visits related to VOC or SCD complications during the 1-year follow-up period were identified. Approximately 85.0% of SCD-related ER visits (n=2,982) or hospitalizations (n=1,449) had VOC as the primary reason for admission; 15.0% of the SCD-related ER visits (n=511 or hospitalizations (n=256) had SCD complications as the primary reason (Fig 5). Of the 256 hospitalizations with an SCD complication as the primary reason, the most common complications were infectious and parasitic diseases (35.5%) and cerebrovascular complications such as seizures (12.5%) and stroke (9.8%). The mean/(SD) and median/(interquartile) for LOS were 7/(7.4) days and 5/(5) days for VOC-related admissions; the mean/(SD) and median/(interquartile) for LOS were 9.7/(11.4) days and 6/(7) days for SCD-complication-related admissions.

**Fig 5.**
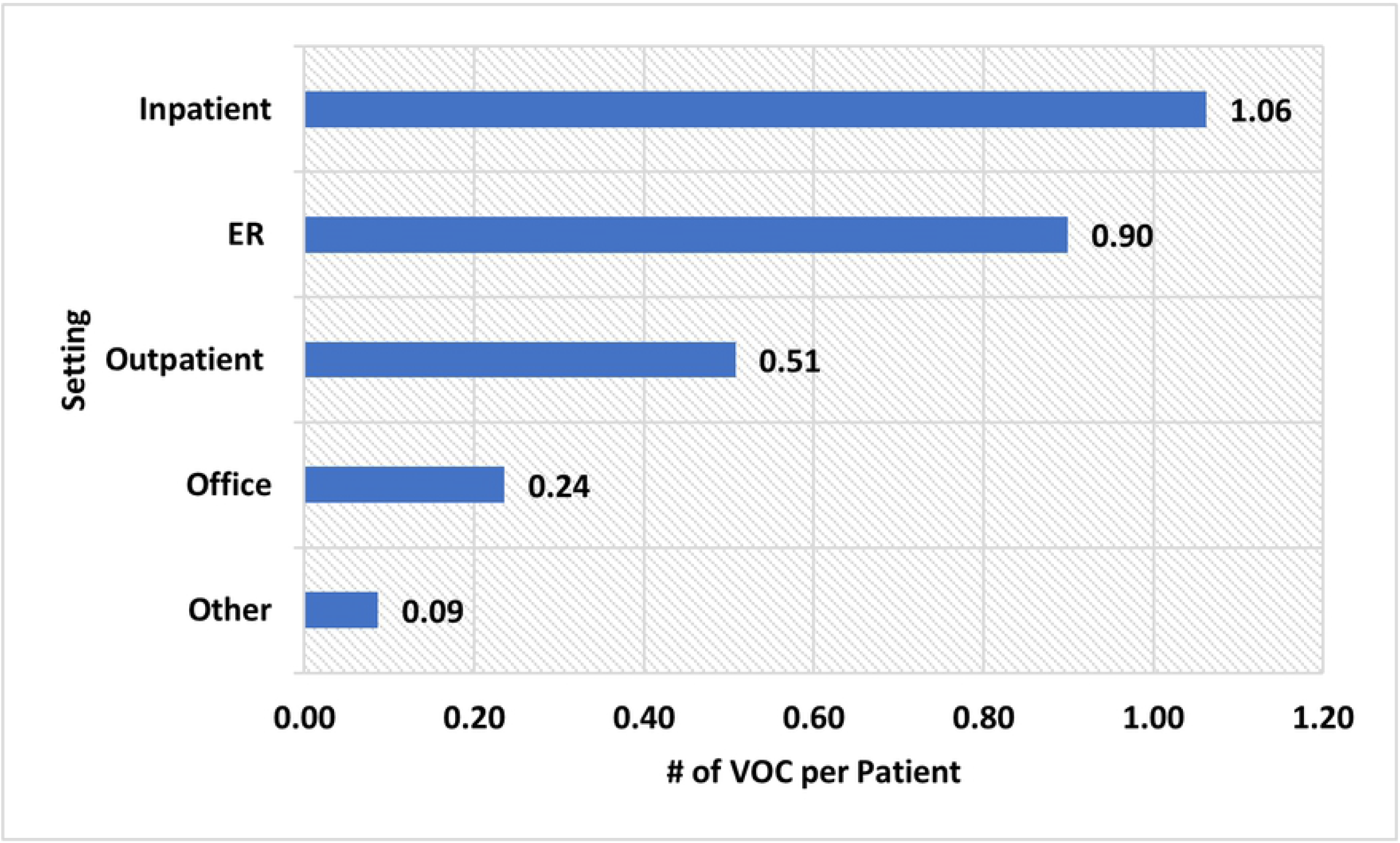
Primary Reason for ER Visits and Inpatient Stay During the First Year of Follow-up. SCD: Sickle cell disease; VOC: Vaso-occlusive crisis.

## Discussion

Using the most recent Medicaid data, this study evaluates the rate of SCD complications in different settings. In addition, the study evaluates the prevalence rate of VOC, the rate of complicated VOC (VOC with concomitant SCD complications) and uncomplicated VOC episodes, and the primary reasons for ER and inpatient admissions.

Over 76,000 VOC episodes were observed during the entire follow-up for the 8,521 adult SCD patients. The median age of the study population was 33 years, indicating that ~50% of the population was aged between 18-33 years. Studies have shown that young patients aged 18-33 years have had the highest mean annual ER visits compared with other age groups [17,23–25]. Studies have found that VOC, which commonly presents as a pain crisis, is the most common reason for hospitalization and ER visits in SCD patients [16,17,26]. Another study showed that >90% of acute hospital admissions of SCD patients were because of severe and unpredictable pain crises [27]. Those findings accord with this study, which observed that VOCs were responsible for 85% of acute medical care for SCD patients, such as ER visits and hospitalization. Moreover, the mean LOS of inpatient visits for SCD or VOC-related admissions was 7 days and 9.7 days for SCD-complication-related admissions, respectively. Similarly, a previous study that examined management approaches for uncomplicated VOC observed an average LOS of 7.8 days for uncomplicated SCD admissions [28].

Studies have shown that VOC episodes in adult SCD patients are actually far more prevalent and severe than what is reported because VOCs are mostly managed at home; this underreporting most likely also indicates an underestimation of VOC prevalence and undertreatment [29]. In the case of this study, discrete VOC episodes within a 3-day gap were recorded as a single episode (eg, multiple ER admissions resulting in hospitalization within a 3-day period would still be calculated as an admission). Per patient we observed an average of 9 episodes requiring health care visits during the entire follow-up period of 2.7 years (ie, an average of about 3.3 VOCs per patient per year). In this study, VOC episodes were more commonly observed in acute settings, such as inpatient and ER, compared to other settings like outpatient.

Moreover, studies have observed infectious disease and pulmonary disease as common complications for SCD patients [16,17]. Those findings accord with this study, which showed that infectious diseases, fever, and pulmonary disorders were the most common concomitant SCD complications observed during the follow-up period.

Although claims data are valuable for the efficient and effective examination of health care outcomes, they are collected for payment and not research. Therefore, certain limitations are associated with their use. For instance, the presence of a diagnosis code on a medical claim does not indicate a positive presence of disease, as the diagnosis code may be incorrectly coded or included as rule-out criteria rather than actual disease.

In addition, this study was conducted on a population that may preclude generalizations to other populations. Specifically, the study setting was within the Medicaid population, which consists of people with disabilities, children of low-income families, pregnant women, parents of Medicaid-eligible children who meet certain low-income requirements, and low-income seniors. These populations are more likely than the general population to have unmet health care service needs.

Some data exclusions were also limitations. We excluded individuals with dual eligibility for both Medicaid and Medicare, for instance, since the data of these observations are not complete. Additionally, due to limited availability, the data of managed care plan patients only included 14 states, and study periods up to 31DEC2013 were the most recent data at the time of the study.

## Conclusions

In a real-world setting, SCD and its related complications result in high acute health care utilization, and VOCs remain the primary reason for SCD patients’ ER visits and inpatient admissions. Further research and interventions are required to facilitate SCD care and VOC management in vulnerable populations.

## Acknowledgements

Editorial support was provided by Michael Moriarty and Chris Haddlesey of STATinMED.

## Supporting information

**S1 Table. List of SCD Complications**. SCD: Sickle cell disease.

